# Structural connectivity gradient associated with a dichotomy reveals the topographic organization of the macaque insular cortex

**DOI:** 10.1101/2022.03.18.484254

**Authors:** Long Cao, Zongchang Du, Yue Cui, Yuanchao Zhang, Yuheng Lu, Baogui Zhang, Yanyan Liu, Xiaoxiao Hou, Xinyi Liu, Luqi Cheng, Kaixin Li, Zhengyi Yang, Lingzhong Fan, Tianzi Jiang

## Abstract

Histology studies revealed that the macaque insular cortex was characterized by the gradual organizations containing agranular, dysgranular and granular insula. However, no consensus has been reached on the elaborate subdivisions of macaque insula. Until now, no neuroimaging study to our knowledge combining connectivity-based gradients and parcellation has been performed to investigate the topographic organization of the macaque insular cortex. In this study, we used high-resolution ex vivo diffusion-weighted imaging data to explore the macaque insular cortex’s global gradient organization and subdivisions. We found a rostrocaudal organization of the dominant gradient in the macaque insula using a diffusion map embedding. Meanwhile, extracting the 25% top and bottom components from the dominant and second gradient, which explained variance over 60% in total within ten gradients, the connectivity-based parcellation method was performed to subdivide each component into two subregions confirmed by the cross-validation analysis. Furthermore, permutations tests identified that two subregions from each component showed significant differences between their connectivity fingerprints. Finally, we found that the dominant and second gradients were significantly correlated with the T1w/T2w and cortical thickness maps in the macaque insula. Taken together, the global gradients combining the subdivisions examined the topographic organization of the macaque insular cortex based on the structural connectivity, which may contribute to a better understanding of the intricate insular cortex anatomy.

## 1. Introduction

Macaque insular cortex, located in the depth of the Sylvian fissure, is structurally and functionally heterogeneous cortical region characterized by the cytoarchitectonic gradients and extensive anatomical connections with the limbic system, prefrontal, orbitofrontal, parietal and temporal lobe. Macaque insula has been demonstrated to be involved in diverse functions, such as emotions, social behavior (Caruana et al. 2011), execution (Di Cesare et al. 2019), reward (Mizuhiki et al. 2012) and also auditory (Remedios et al. 2009), gustatory (Yaxley et al. 1990), sensorimotor, orofacial motor (Jezzini et al. 2012), etc. A unique characteristic of the insula is the presence of von Economo neuron (VEN), which is associated with self-awareness and social cognition, and was originally observed only in the great ape and humans, but not other primates. However, a recent architectonic study demonstrated that the VENs also existed in the anterior insula of the macaque (Evrard et al. 2012), suggesting that the insula of macaque monkeys plays a critical role in evolutional and comparative neuroscience (Evrard 2019).

Structural organization of macaque insula is primarily obtained from cyto-, or myelo-, and receptor-architectonic studies. Using Nissl and myelin staining, Mesulam and Mufson proposed the concept of granularity and subdivided macaque insula into three architectonic areas, including argranular, dysgranular and granular insula (Ia, Id and Ig), which were just delineated by the circular sulcus and limen insulae (Mesulam and Mufson 1982). Subsequently, Carmichael and Price used multiple stains to subdivide macaque orbital and medial prefrontal cortex and found that the anterior insula subregion Ia was extended onto the posterior orbital surface and could be subdivided into five subregions, including medial, intermediate, lateral, posteromedial and posterolateral insula (Iam, Iai, Ial, Iapm and Iapl) (Carmichael and Price 1994a). Recently, using multiple immunohistochemical stains, Gallay et al. revealed that macaque insula could be into eight subregions (Ia1-2, Id1-3, Ig1-2 and G) with parallel ventral to dorsal gradients by the cytoarchitectonic features of layer II-V (Gallay et al. 2012a). After confirming the VENs in the Ia of macaque insula, Evrard et al. employed Nissl and Myelin techniques to subdivide macaque insular cortex into fifteen subregions (Evrard et al. 2014), which showed medial-lateral or ventrodorsal pattern along the rostrocaudal axis of macaque insula territory. Therefore, microstructural evidence has demonstrated that the macaque insular cortex displays anterior-middle-posterior divisional patterns at a coarse level. In contrast, the highly specialized subregions of macaque insula remain not well-established.

Following the development of neuroimaging techniques, accumulating studies have focused on the organization of the human insular cortex. Initially, most neuroimaging studies performed connectivity-based methods to parcellate human insula with clear boundaries. However, recent neuroimaging research revealed that human insula represented global gradients in macroscopic and microscopic organizations. For example, based on probabilistic tractography, Cerliani et al. utilized Laplacian eigenmaps to explore the organization of human insula, and unveiled the rostrocaudal gradients of structural connectivity (SC) variation across the insula territory (Cerliani et al. 2012b). More recently, in one fMRI study, Tian et al. revealed that human insula could be characterized as a continuum of gradual change along the rostrocaudal axis and parcellated human insula into two subregions with anterior-posterior pattern (Tian and Zalesky 2018). Using multi-shell diffusion-weighted imaging (DWI), Menon et al. used RTOP representing granularity to investigate the microstructure of the human insular cortex and found gradients along the anterior-posterior and dorsal-ventral axes (Menon et al. 2020). Using T1w/T2w maps, Royer et al. uncovered two principal gradients of human insula myeloarchitecture showing one from ventral anterior to the posterior insula and the other from dorsal anterior to both ventral anterior and posterior insula (Royer et al. 2020). Taken together, apart from neuroimaging research with connectivity-based parcellation method on insula organization, several studies also considered that the insular cortex was characterized with the overall gradual change along the rostrocaudal or ventrodorsal axis. Until now, no neuroimaging study to our knowledge used connectivity-based parcellation or gradient method or both to explore the macrostructure of macaque insular cortex specifically. Therefore, the topographic organization of the macaque insula, representing the connectivity variation, remains unknown.

In this study, we used high spatial and angular resolution ex vivo macaque DWI data to investigate the topographic organization of the macaque insular cortex. First, we used a diffusion map embedding to examine macaque insula global gradients. Second, extracting the top and bottom components from the first two gradients, we performed connectivity-based parcellation to subdivide each component to uncover the elaborated macaque insula organization. Third, using connectivity fingerprints, we examined the structural connectivity differences between the subregions subdivided from one component. Finally, we explored the relationships between macaque insula global gradients and two neuroimaging indices, including T1w/T2w and thickness maps. Here, we hypothesized that the globally spatial representation of macaque insula would show a rostrocaudal gradient, and the elaborated subdivisions of macaque insula would show medial-lateral or ventrodorsal pattern along the rostrocaudal gradient.

## 2. Materials and methods

### 2.1 Data acquisition

#### 2.1.1 Ex vivo macaque brains

Eight ex vivo macaque (*Macaca mulatta*; 2 males, 6 females; 5.6 ± 1.06 years of age) brains were collected, and the experimental protocols were approved by the National Animal Research Authority of China and the Ethics Review Committee of Biomedical Research of the Institute of Automation, Chinese Academy of Sciences. MRI data contained T2-weighted (T2w) and DWI images which were obtained from a 9.4-T horizontal animal MRI system (Bruker BioSpec 94/30 USR) with Paravision 6.0.1. The gradients were equipped with a slew rate of 4570 mT/m/ms and maximum strength of 660 mT/m. A 72 mm inner diameter quadrature radiofrequency coil achieved the radiofrequency transmission and reception. Prior to imaging, each postmortem macaque brain was soaked in 1L phosphate buffered saline with 1mM/L gadopentetate dimeglumine (Gd-DTPA) at 4 °C and shook daily for at least 1 month to reduce T1 relaxation time (D’Arceuil et al. 2007). A 3D spin-echo EPI diffusion weighted sequence was used to acquire multi-shell DWI with echo time (TE) = 25 ms, repetition time (TR) = 200 ms, 186 or 129 diffusion directions (7 brain DWI images, 6 b = 0 s/mm^2^, 60 b = 1200 s/mm^2^, 120 b = 4800 s/mm^2^; 1 brain DWI image, 6 b = 0 s/mm^2^, 30 b = 1200 s/mm^2^, 93 b = 4800 s/mm^2^), field of view (FOV) = 66.6 × 54.0 × 72.0 mm^3^, flip angle (FA) = 90°, matrix = 148 ×120 × 160 and voxel size = 0.45 × 0.45 × 0.45 mm^3^ without gap. T2w images were obtained by a 3D MSME sequence with TE = 36 ms, TR = 2000 ms, FOV = 72.0 × 54.0 × 80.1 mm^3^, FA = 90°, matrix = 240 × 180 × 267 and voxel size = 0.3 × 0.3 × 0.3 mm^3^.

#### 2.1.2 In vivo macaque brains

Eight in vivo macaque (*Macaca mulatta*; male) brain MRI data used in this study was obtained from TheVirtualBrain (Shen et al. 2019). The Animal Use Subcommittee approved the experimental protocols of the University of Western Ontario Council on Animal Care. They were following Canadian Council of Animal Care guidelines. The neuroimaging protocol included T1-weighted (T1w) and DWI images acquired by the 7.0-T Siemens MAGNETOM head scanner with a gradient (Siemens AC84 II, Gmax = 80 mT/m, SlewRate = 400 T/m/s). T1w images were collected by an MP2RAGE acquisition with TR = 6500 ms, TE = 3.15 ms, TI1 = 800 ms, TI2 = 2700 ms, matrix = 256 × 256, 128 slices and voxel size = 0.5 × 0.5 × 0.5 mm^3^, and only T1w images were analyzed in our study.

### 2.2 Data preprocessing

Preprocessing of macaque ex vivo DWI data included reorienting the initial brain space into RAS space, denoising (Veraart et al. 2016), removing ringing artifacts (Kellner et al. 2016), eddy currents correction (Graham et al. 2016), brain-tissue extraction on b0 image (Smith 2002) and refining brain mask manually, which were respectively implemented in the Medical Image Processing, Analysis and Visualization (MIPAV) (https://mipav.cit.nih.gov/), MRtrix3 (Tournier et al. 2019), FMRIB Software Library (FSL v6.0) (Woolrich et al. 2009; Jenkinson et al. 2012) and ITK-SNAP (http://www.itksnap.org/). N4BiasFieldCorrection (Tustison et al. 2010) within Advanced Normalization Tools (ANTs) (http://stnava.github.io/ANTs/) was performed to correct a b0 image for the subsequent registration. Then DTIFIT (Smith et al. 2004) was used to fit a diffusion model to check the principal eigenvector within brain voxels in the native space. After confirming the principal eigenvector, BEDPOSTX (Jbabdi et al. 2012) was performed to estimate the distribution of three fiber orientations at each voxel.

Given the use of Gd-DTPA and the absence of ex vivo T1w image, we inverted the contrast of T2w or b0 image to obtain a fake T1w image (Ambrosen et al. 2020) which was then preprocessed by the HCP-NHP pipeline (Autio et al. 2020), including warping the individual volume image into the standard Yerks19 template (Donahue et al. 2018) using FSL, surface construction using Freesurfer (https://surfer.nmr.mgh.harvard.edu/) and mapping the individual space into the Yerk19 surface using the multimodal surface matching algorithm (Robinson et al. 2014) (details in Fig. S1).

The T1w images of macaque in vivo TheVirtualBrain dataset were also preprocessed by the HCP-NHP pipeline (details in Fig. S1).

### 2.3 Definition of macaque insula ROI

Our macaque insula ROI, mainly defined by the sulcus and gyrus, was delineated in a postmortem macaque b0 template (CIVM) (Calabrese et al. 2015a) with labels of Paxinos et al.’s macaque atlas (Paxinos et al. 2009) at the 0.3-mm isotropic resolution. More specifically, first, we extracted the insula mask, including Ia, Id, Ig, and IPro (insular proisocortex), which was defined in the CIVM atlas. This insula mask’s ventral and dorsal extent was bounded by the inferior and superior limiting sulcus of the insula, respectively, and its anterior extent bordered ventrally on the piriform cortex. Second, we delineated the most anterior insula mask with five regions, including Iam, Ial, Iapm, Iai and Iapl, which were defined in the D99 template (Reveley et al. 2016) with labels of Saleem et al.’s macaque atlas (Saleem and Logothetis 2007). These five regions were transformed from the D99 space into the CIVM space using ANTs. In the CIVM space, the lateral extent of this mask was delimited by the inferior limiting of the insula and its extended line, and the medial extent bordered posteriorly on the anterior olfactory nucleus and medially piriform cortex. Finally, we merged these two masks into our macaque insula ROI in the CIVM space, and the detailed volume ROI was shown in Fig. S2.

### 2.4 Macaque insula SC gradients

We aimed to construct a macaque insula connectome in the common volume space, a symmetrical cross-correlation matrix, which could reveal the similarity of connectivity patterns within voxels, and then be unveiled by the diffusion embedding to explore global insula spatial distributions. For each ex vivo macaque brain, nonlinear registration (Avants et al. 2014) within ANTs was applied to estimate a deformation field between the individual b0 image and CIVM b0 template with an isotropic resolution of 0.3 mm. Given a decrease of the voxel number transformed from the high-resolution b0 template to a low-resolution individual b0 image, we resampled the macaque insula template mask into the 0.6-mm isotropic resolution, and then labeled all the voxels within the insula mask from 1 to M. According to the estimated deformation field, the insula mask was warped from the CIVM space with the isotropic resolution of 0.6 mm into the individual space with a 0.45-mm isotropic resolution using nearest-neighbor interpolation. In the individual space, the probability between each voxel in the insula mask and the brain voxels (N) was calculated by the probabilistic tractography using Probtrackx2 (Behrens et al. 2007) with the step length of 0.25 mm (Bryant et al. 2020) and curvature threshold of 0.2, and the pial surfaces generated in the previous HCP-NHP pipeline were treated as a stop mask to prevent fiber tracking from crossing sulci. Five thousand streamlines per insula voxel were tracked to calculate the connectivity profiles, and we averaged the connectivity profiles of voxels with the same label one by one to obtain the individual M-by-N matrix. Then, a cross-correlation M-by-M matrix was calculated to reveal the similarity between the voxels with different labels. Finally, we averaged all the subjects cross-correlation matrices to obtain a group-level cross-correlation M-by-M matrix with the min-max normalization.

A diffusion map algorithm (Berry and Harlim 2016) was applied to the group-level cross-correlation matrix to obtain ten gradients of insula. The variance was calculated within each gradient and normalized to acquire the explained variance of the gradient. The gradients in the CIVM space were warped into the Yerks19 space using nonlinear registration within ANTs. We normalized each gradient in the volume space and then mapped it to the 32k Yerks19 surface for display using Connectome Workbench (https://www.humanconnectome.org/).

### 2.5 SC-based parcellation

In the CIVM space, we extracted the top and bottom 25% of all insula voxels in the dominant gradient, and also extracted the top and bottom 25% of the rest insula voxels in the second gradient. The four components were treated as the ROIs, which were subdivided using the SC-based parcellation frame (Fan et al. 2016). The main steps can be summarized as follows: first, the ROI was warped from the CIVM space into the individual space using the estimated deformation field. Second, individual pial surfaces were conducted as the stopping mask, and 5000 streamlines per voxel in the individual ROI were tracked to construct the connectivity matrix between the ROI and brain voxels using Probtrackx2. Third, in the individual space, we obtained the cross-correlation matrix by multiplying the connectivity matrix by its transpose, which was used as an input to the spectral clustering algorithm (Ng et al. 2002). Fourth, cluster number was set from 2 to 6 and the individual parcellation results were warped into the CIVM space. Finally, for each cluster solution, the maximum probabilistic map of one subregion was acquired by identifying that the probability of voxel belonging to the subregion is more than 50% across the subjects. Additionally, to avoid the arbitrary selection of clusters, the cross-validation indices Cramer’s V and topological distance (TpD) (Li et al. 2017) were calculated to choose the optimal cluster across all subjects.

### 2.6 SC pattern

After obtaining four components from the insula gradients and subregions from the subdivision on the component, we characterized their connectivity profiles using the probabilistic tracking strategy used in the SC parcellation. For the component or subregion, 50000 samples per voxel were tracked to the whole brain using Probtrackx2, and also the pial surfaces were served as the stop mask. For each subject, the connectivity profile of one component or subregion was only focused on the ipsilateral hemisphere and transformed into the CIVM space. To further reduce the false positive connections, 3.08 × 10^−5^ for the samples (He et al. 2020) was used to binarize the connectivity profile. We averaged connectivity profiles of the subregion across the subjects to get the probabilistic connectivity profile, and the threshold 0.5 was used to obtain the group-level connectivity profile for the given subregion.

Meanwhile, we also performed connectivity fingerprints (Passingham et al. 2002) to reveal the connectivity pattern between insula component or subregion and ipsilateral target ROIs. The main procedures were summarized as follows: first, all cortical and subcortical regions in the CIVM atlas with Paxinos et al.’s labels of macaque atlas (Paxinos et al. 2009) were extracted as the target ROIs. Second, after obtaining the connectivity profiles thresholded in the CIVM space, the connectivity values were calculated by averaging the voxel values within all the target ROIs for each component or subregion, and then we averaged the connectivity values across the subjects to obtain the initial connectivity fingerprints. Third, for all components or subregions, we thresholded the connectivity fingerprints at >0.001 and then merged some regions with similar connectivity patterns into one homogenous region. For example, area 10 and its dorsal, medial, ventral part were merged into one target ROI. Thus, a target ROI family was generated, containing 29 cortical and 6 subcortical regions (details in Tabel S1). Fourth, in the one subject, we recalculated the connectivity fingerprints using the new target ROIs family again for all the components or subregions with the previous step 2. The connectivity fingerprints of one subject were normalized by summing all the connections to 1 (Mars et al. 2012; Xia et al. 2017) within one component or subregion in all target ROIs and subsequently within all the components or subregions in one target ROI. Finally, the connectivity fingerprints of the components or subregions were constructed by averaging the individual normalized connectivity fingerprints across the subjects.

A permutation test (Mars et al. 2016) was performed to investigate the difference of connectivity fingerprints between the two subregions that originated from the component subdivisions. More specifically, after obtaining the normalized fingerprints of all subjects, we calculated the Manhattan distance between the connectivity fingerprints of the two subregions with 35 targets ROIs within all subjects to characterize the observed difference between the connectivity patterns of the two subregions. The hypothesis is that the difference between the connectivity patterns of the two subregions was higher than expected by chance. Subsequently, the Manhattan distances between the subregions were calculated for the 10000 permutations for connectivity fingerprints of two subregions. If the observed Manhattan distance was higher than the criterion derived from the permutation test (*p* < 0.05), the two subregions subdivided from the component would show significant differences in the connectivity pattern.

### 2.7 Relation to the T1w/T2w and cortical thickness map

The cortical thickness maps in the Yerks19 space (Donahue et al. 2018) were generated by the HCP-NHP pipeline in the macaque in vivo dataset. By averaging the cortical thickness maps across the subjects, we obtained the group-level cortical thickness map. The correlation analysis was performed to investigate the relationship between the insula dominant gradients and the insula cortical thickness maps by the vertex values of the insula in the Yerks19 space. To explore the relationship between insula SC gradients and T1w/T2w map, we used the standard T1w/T2w map in the HCP-NHP pipeline, and then separated and resampled it into the 32k space using Connectome Workbench. Likewise, the correlation analysis was performed to investigate the relationship between the insula dominant gradients and T1w/T2w maps.

## 3. Results

### 3.1 SC gradients of macaque insula

Based on the macaque insula SC connectome, diffusion maps revealed global spatial distributions of the insular cortex where the top two insula gradients explained the variance over 50% in total. We only focused on the top two gradients, and the others explained the variance all below 10%. The dominant gradient (G1) explained 45.7% and 41.5% variance of the left and right insula respectively (Fig. 1A). It showed a gradual increase along the rostrocaudal axis (Fig. 1B, top). The second gradient (G2), accounting for 16.0% and 15.1% variance for the left and right insula (Fig. 1A), showed the highest proportion in the middle insula and a gradual decrease from the middle to the rostral and caudal direction separately (Fig. 1B, bottom).

**Figure 1.**
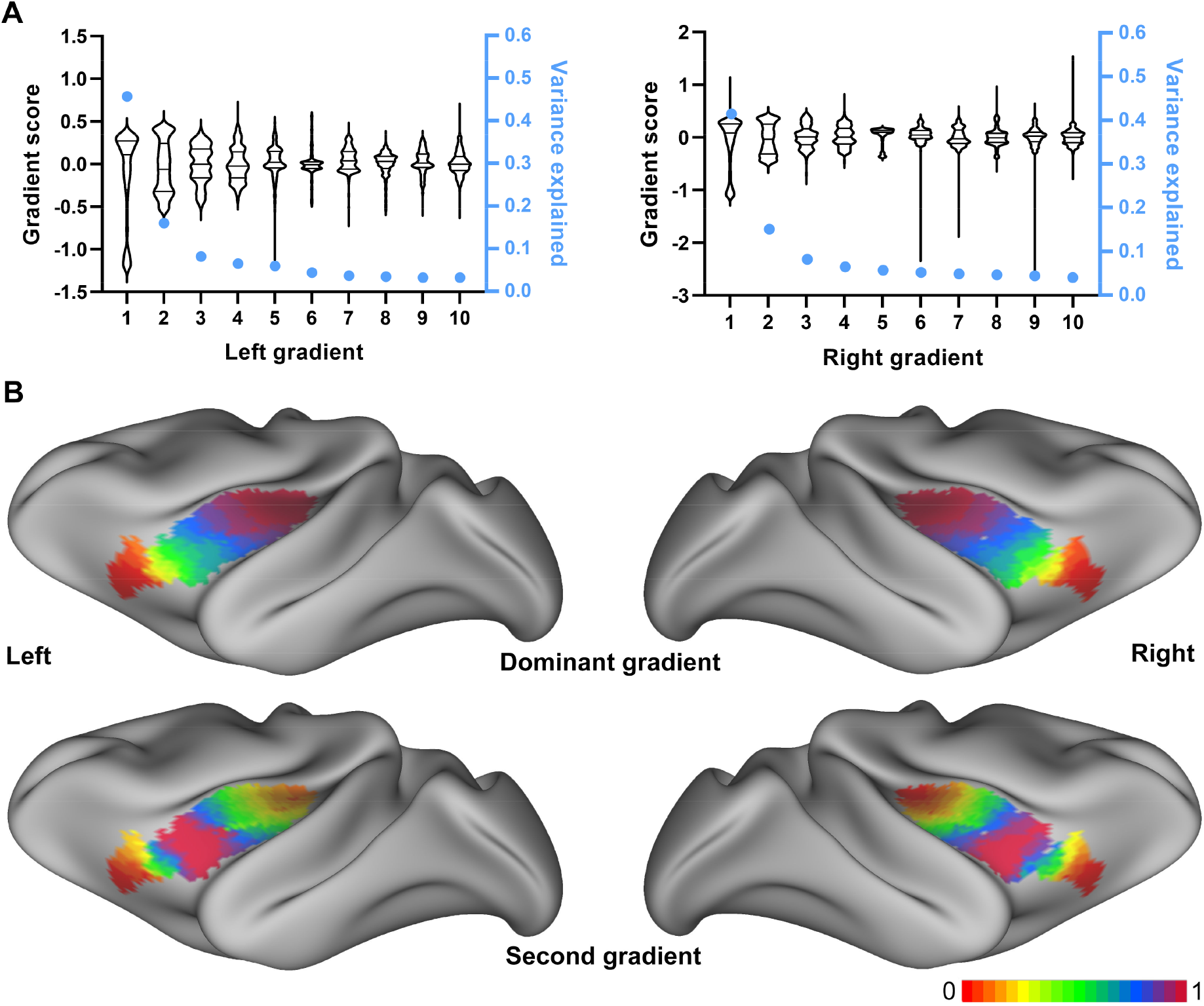
Macaque insula gradients based on structural connectivity. (A) Ten insula gradients distribution and their variance explained. (B) The top two insula gradients mapped into Yerks19 surface space. The gradients were normalized to the maximum value in the range of 0 to 1 for visualization.

### 3.2 Parcellation of macaque insula based on SC gradients

The four components (INS1-4), including the top and bottom 25% of G1 and the additional top and bottom 25% of G2, were extracted and merged into the initial partition of the insula with the parallel rostral to the caudal pattern (Fig. 2A). With the highest CV and TpD indices, these four components from INS1 to INS4 were all subdivided into two subregions (Fig. 2B), including anteromedial and anterolateral Ia (Iaam and Iaal), posterodorsal and posteroventral Ia (Iapd and Iapv), dorsal and ventral Id (Idd and Idv), and dorsal and ventral Ig (Igd and Igv) (Fig. 2C).

**Figure 2.**
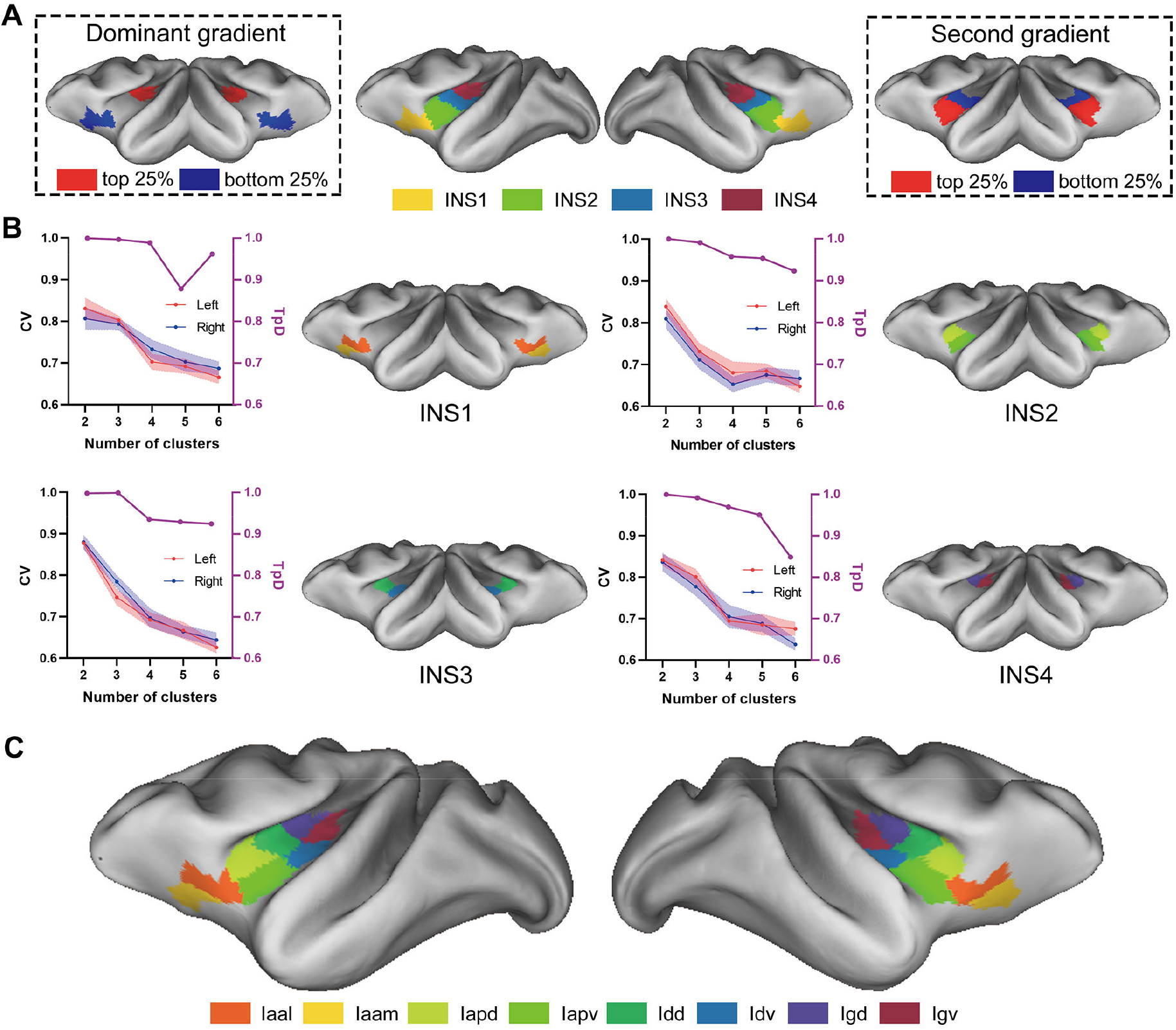
Subdivisions of macaque insula. (A) Initial insula partition merged from the dominant and second gradient with top and bottom 25% portion. (B) Validation indices and subdivisions in 2 clusters for all four components. Cramer’s V (CV) described consistency of parcellation across subjects, and topological distance (TpD) measured the symmetry of parcellations between hemispheres. Higher values of CV and TpD indicated good consistency and symmetry, respectively. (C) The final insula subdivision with 8 subregions.

### 3.3 SC fingerprint of macaque insula subregion

INS1, adjacent to area 13 medially, was located in the rostral part of the macaque insula, and was subdivided into two subregions, including the medial portion Iaam and the lateral portion Iaal (Fig. 2B). INS1 had strong ipsilateral connections with area 10, 46, 47, 11, 13, 14, 25, 32, 36R, TP and subcortical regions Acb, Str, Amyg, Pir (Fig. 3A). Using permutation tests, the two subregions showed significant differences (LH: *p* < 0.0001, RH: *p* < 0.0001) in the connectivity fingerprints and Iaam showed higher connections with area 14, 25, 36R, and Pir, whereas area 10, 46, 47, 11, 13, 32, Tha, Acb and Amyg had higher connections with Iaal (Fig. 4A).

**Figure 3.**
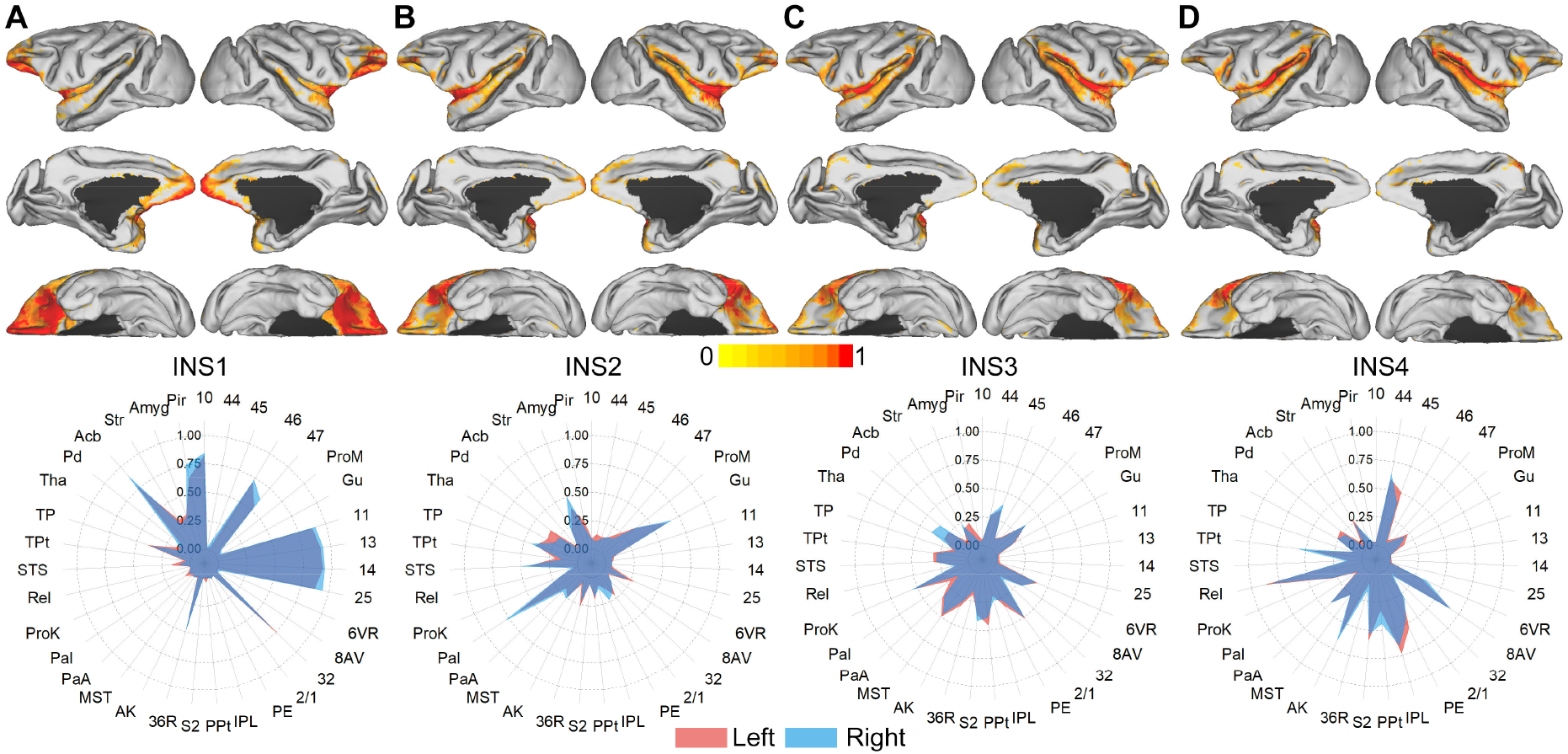
The connectivity patterns of four components, including INS1 (A), INS2 (B), INS3 (C), and INS4 (D). The connectivity profile was mapped into the F99 surface space (Van Essen 2002) for visualization using Caret5 and focused on the ipsilateral hemisphere.

**Figure 4.**
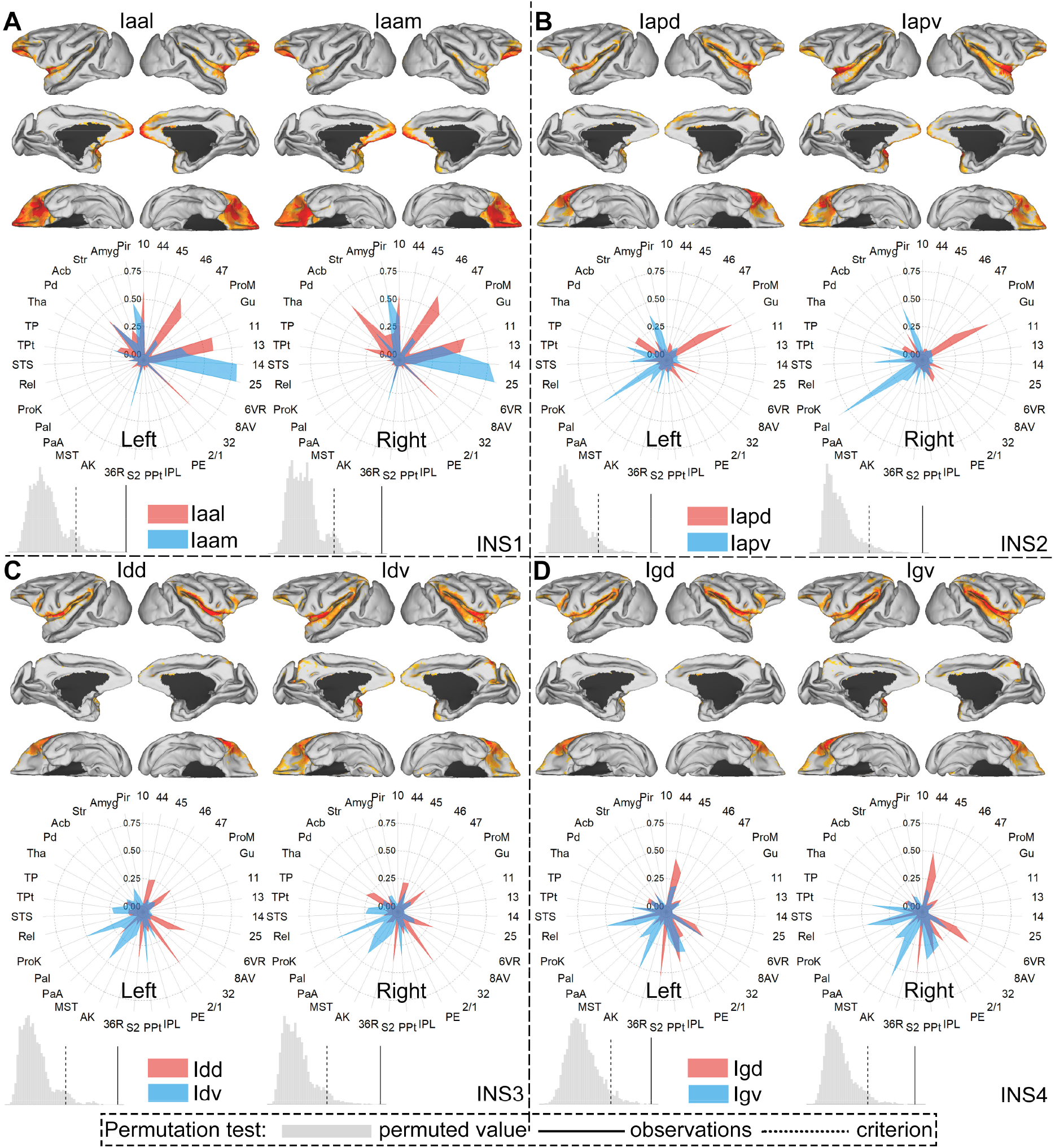
Connectivity patterns of each two subregions subdivided from four components INS1 (A), INS2 (B), INS3 (C), and INS4 (D). Connectivity profiles focused on the ipsilateral hemisphere for one subregion. Permutation tests (histograms) were used to identify differences between connectivity fingerprints of two subregions. The observations, the Manhattan distance between two fingerprints, was higher than the criterion of permutations, indicating significant differences between the connectivity fingerprints of two subregions.

INS2 covered the anterior domain of the middle insula just posterior to INS1, which was parcellated into two subregions, including the dorsal portion Iapd and the ventral portion Iapv (Fig. 2B). INS2 had high ipsilateral connections with area ProM, Gu, 25, PaI, STS, TP and subcortical regions Tha, Pir, Str, Pd (Fig. 3B). Significant differences (LH: *p* < 0.0001, RH: *p* < 0.0001) were also found by permutation tests between the connectivity fingerprints of these two subregions. Iapd had higher connections with area ProM, Gu, 6VR, 2/1, Tha, and Pd, whereas area PPt, 36R, MST, PaA, PaI, STS, TP and Amyg showed higher connections with Iapv (Fig. 4B).

INS3 occupied the posterior domain of the middle insula just posterior to INS2, and was subdivided into two subregions, including the dorsal portion Idd and the ventral portion Idv (Fig. 2B). INS3 had strong ipsilateral connections with areas 44, 45, ProM, Gu, 6VR, 8AV, 2/1, PE, PPt, S2, AK, MST, PaA, ProK, STS, TPt, and subcortical regions Tha, Pd (Fig. 3C). Permutations tests identified significant differences (LH: *p* < 0.0001, RH: *p* = 2.00 × 10^−4^) between connectivity fingerprints of these two subregions. Idd showed higher connections with areas 44, 45, ProM, Gu, 6VR, 2/1, S2, Tha and Pd, whereas Idv had higher connections with 36R, AK, MST, ProK, STS, and TPt (Fig. 4C).

INS4, adjacent to the S2 and ProK laterally, was located in the posterior domain of the insula and was parcellated into two subregions, including the dorsal portion Igd and the ventral portion Igv (Fig. 2B). INS4 had strong ipsilateral connections with areas 44, 45, 6VR, 8AV, PE, IPL, PPt, S2, AK, MST, PaA, ProK, ReI, TPt, and subcortical regions Tha, Pd (Fig. 3D). Significant differences (LH: *p* < 0.0001, RH: *p* < 0.0001) were identified by permutation tests between the connectivity fingerprints of these two subregions. Igd showed higher connections with areas 44, 45, 6VR, 8AV, and S2, whereas area PE, PPt, AK, MST, PaA, ProK and ReI had higher connections with Igv (Fig. 4D).

### 3.4 Relationships between gradients and T1w/T2w, cortical thickness maps

Macaque insula G1 correlated positively with insula T1w/T2w map (LH: *r* = 0.46, *p* < 0.0001; RH: *r* = 0.43, *p* < 0.0001) (Fig. 5A), and negatively with insula thickness map (LH: *r* = −0.31, *p* < 0.0001; RH: *r* = −0.28, *p* < 0.0001) (Fig. 5B). However, G2 correlated negatively with insula T1w/T2w map (LH: *r* = −0.43, *p* < 0.0001; RH: *r* = −0.40, *p* < 0.0001) (Fig. 5A), and positively with insula thickness map (LH: *r* = 0.26, *p* < 1.16e-14; RH: *r* = 0.29, *p* < 6.31e-19) (Fig. 5B).

**Figure 5.**
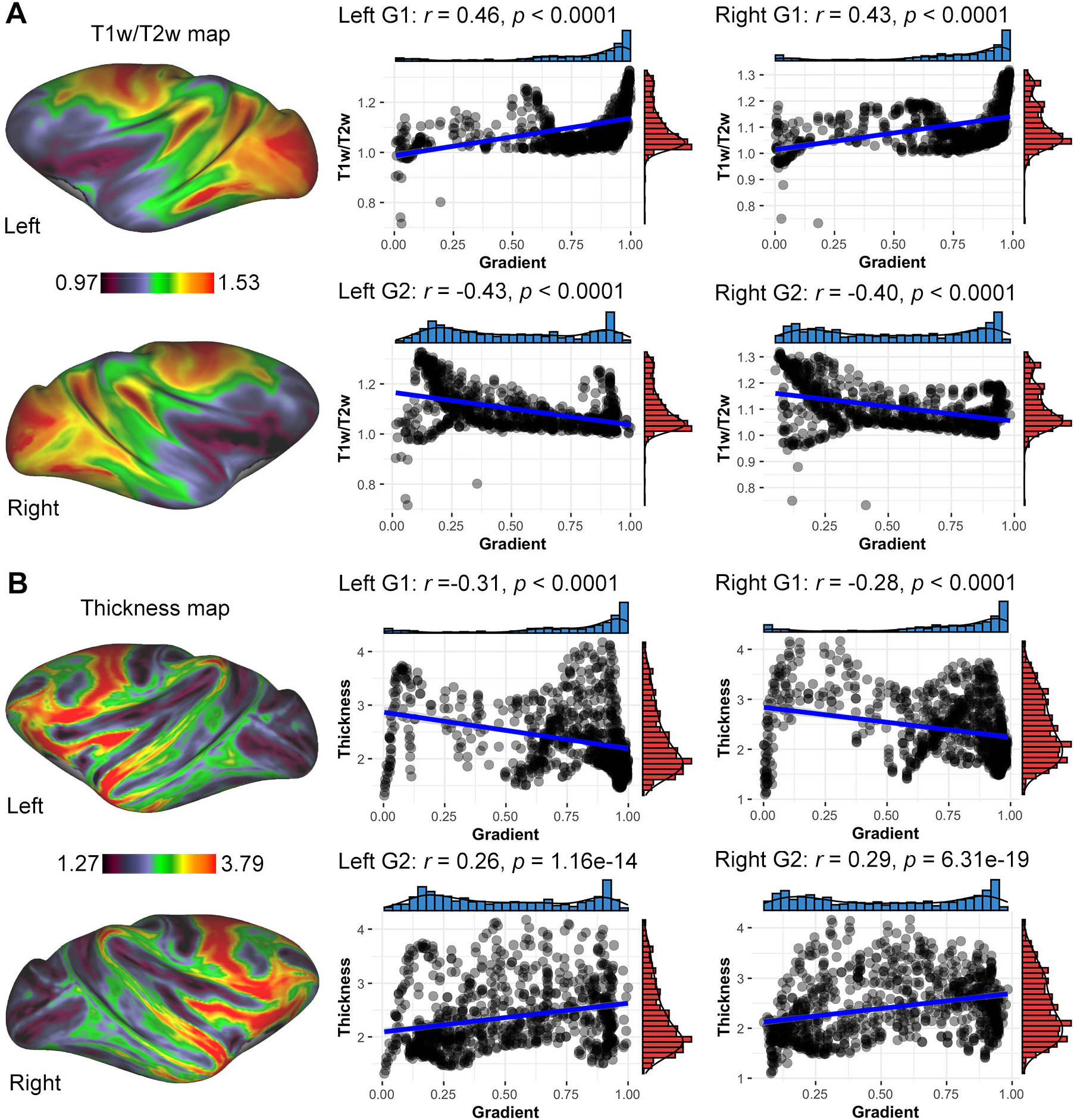
Insula gradients captured insula microstructure and morphology. (A) Insula gradients revealed the significant correlations with T1w/T2w map. (B) Insula gradients were associated with cortical thickness.

## 4. Discussion

Based on structural connectivity, this study investigated global gradients and subdivisions in the macaque insular cortex and also explored the relationship between insula global gradients and its morphology and microstructure. The insula G1 revealed the gradually rostrocaudal increases, whereas the insula G2 represented gradual increases from the rostral and caudal portions to the middle domain. Extracting four components from the first two insula gradients, a connectivity-based parcellation method was performed to subdivide each component into two subregions, showing medial-lateral or dorsal-ventral parcellation pattern. Moreover, these two subregions showed significant differences between their connectivity fingerprints by permutation tests. In addition, insula gradients also captured the characteristics of T1w/T2w and thickness maps. Overall, macaque insula gradients or subdivisions using neuroimaging provided insights to comparative or translational medicine research.

### 4.1 Macaque insula gradients and parcellations

We examined the spatial representation of structural connectivity variation across macaque insula territory and found a rostrocaudal gradient (Fig. 6A top). This result was similar to a cytoarchitectonic study with multiple stains reporting that parallel ventral to dorsal gradients were found in the middle layer of the macaque insula (Fig. 6B) (Gallay et al. 2012b). Our result was consistent with a human neuroimaging study showing that probabilistic tractography uncovered a rostrocaudal trajectory of connectivity variability in the human insula (Cerliani et al. 2012a). However, the comparability across species is not clear and requires further examination. Besides, we constituted a 4-subregion parcellation pattern by extracting the top and bottom 25% components from the first two gradients instead of directly hard clustering algorithms to subdivide macaque insula. Given the unstable boundaries in the middle insula by the hard clustering reported in a previous study (Nanetti et al. 2009), such extract method making up 4-subregion pattern could guarantee that the two subregions from one insula gradient represented the maximum connectivity variations. More interestingly, the three subregions INS2-4 posterior to the limen insula in the 4-subregion pattern spatially arranged by a cytoarchitectonic study of the macaque insula with an anteroposterior parcellation (Fig. 6C) (Calabrese et al. 2015a; Paxinos et al. 2009). Moreover, the subdivision of INS2, which was parcellated into Iapd and Iapv, was similar to the Ia and IPro in the Paxinos et al.’s macaque atlas that Ia showed more SMI reactivity than IPro in the infragranular layers (Paxinos et al. 2009). In case of these three subregions, the subdivision results all revealed the ventrodorsal pattern, similar to the cytoarchitectonic study showing the dorsal and ventral architectonic areas of the macaque posterior insula (Fig. 6D) (Evrard et al. 2014). Concerning the subregion INS1 in the posterior surface of the orbitofrontal lobe, the subdivision of this region showed a medial-lateral pattern which also corresponded to a previous histology study subdividing the proportion of insula anterior to limen insula into Iam, Iai, Ial, Iapm, and Iapl (Carmichael and Price 1994b). Taken together, our results revealed the structural organization of macaque insula, showing from the global anteroposterior gradient pattern to the mediolateral or ventrodorsal subdivision pattern locally, which were both consistent with the previous cytoarchitectonic studies on macaque insula.

**Figure 6.**
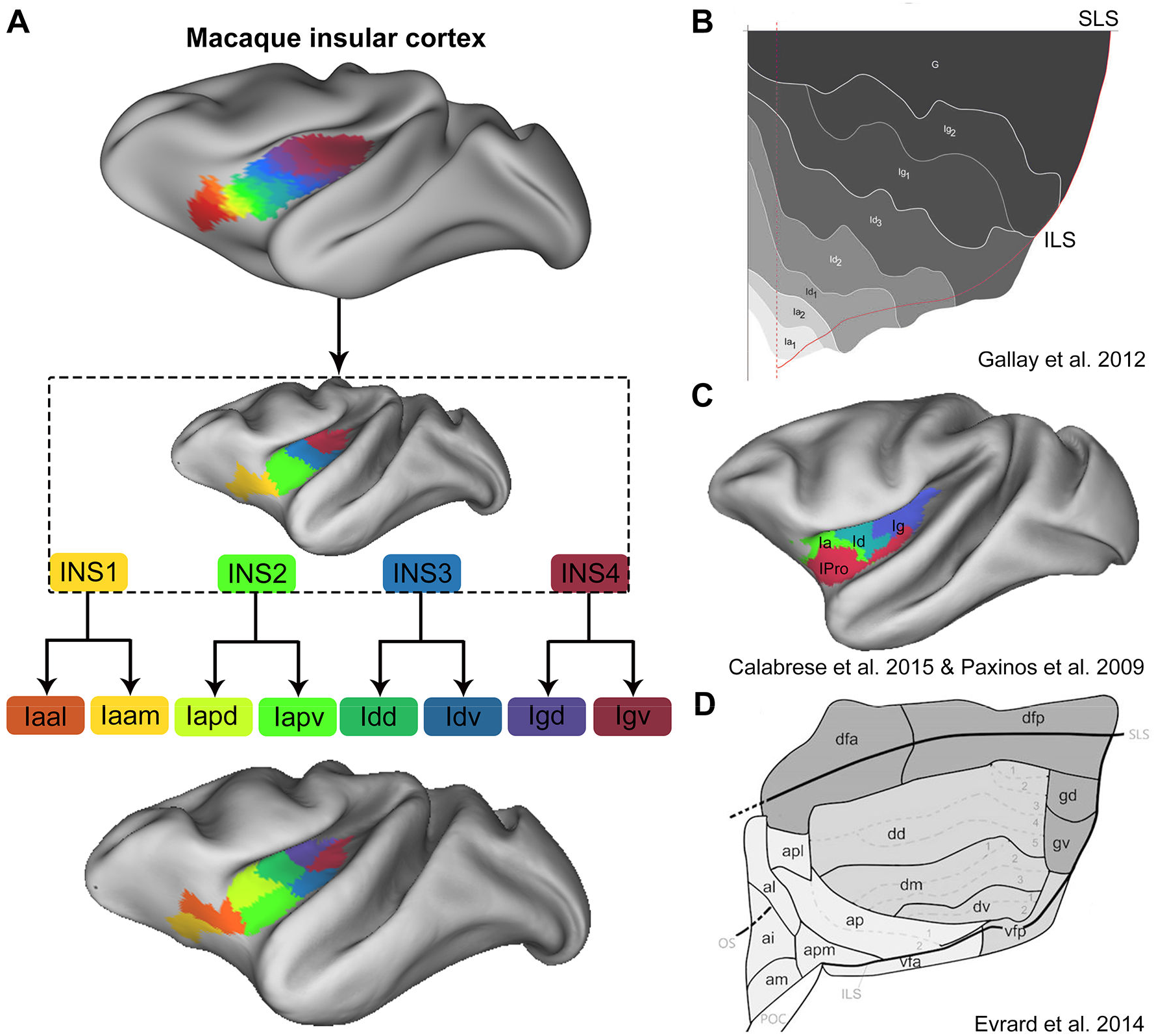
Comparison between macaque insula topographic organization in our study and the prior insula histological architecture. (A) Insula topographic organization. From top to bottom, macaque insula was obtained by structural connectivity gradients, 25% components extraction, and connectivity-based subdivisions of all components sequentially. (B) Macaque insula cytoarchitecture gradients were obtained from Gallay et al.’s study (Gallay et al. 2012b). (C) The insula cytoarchitecture of Paxinos et al.’s (Paxinos et al. 2009) and Calabrese et al.’s (Calabrese et al. 2015b) macaque atlas was mapped to the F99 surface space. (D) Macaque insula cytoarchitecture was obtained from Evrard et al.’s study (Evrard et al. 2014)

### 4.2 Macaque insula SC

Apart from research on the structural organization of macaque insula, the present study also investigated the structural connectivity of the insula subregions. INS1 had strong structural connectivity with the lateral prefrontal cortex, orbitofrontal cortex, anterior cingulate gyrus, perirhinal cortex, Pir, and Acb. More specifically, the lateral subregion Iaal, subdivided from the INS1, showed higher connections with areas 10, 46, 47, 32, 11, 13, and Acb, whereas the medial subregion Iaam connected strongly with area 14, 25, and Pir. The previous tracer study reported anatomical connections between area 10 and the posterior orbitofrontal lobe (Petrides and Pandya 2007). The macaque frontal pole was thought to play a critical role in cognitive processing, such as complex and higher-order behaviors (Boschin et al. 2015). Previous lesion studies (Murphy and Bachevalier 2020) and tasks (Setogawa et al. 2019) revealed that orbitofrontal areas 11, 47, and 13 played vital roles in the attention, reward and may also participate in social behavior. Thus, the connections between Iaal and these regions may indicate that the lateral subregion of INS1 was associated with the higher cognitive functions. In addition, in one tracer study (Carmichael et al. 1994), Pir was connected with two cytoarchitectonic insula areas (Iam and Iapm) corresponding to the subregion Iaam in our research. Pir plays a vital role in the processing and encoding of olfactory information (Boyett-Anderson et al. 2003), and area 14 also participants in olfactory-related function (Ongur and Price 2000). Therefore, the medial subregion Iaam of INS1 may play an essential role in the processing of olfactory information. For areas 25 and 32, the recent study suggested that these two regions of the primate were involved in the opposite roles in regulating negative emotion, such as the cardiovascular and behavioral correlates (Wallis et al. 2017). In our study, the two subregions, Iaam and Iaal, showed high connections, respectively, with areas 25 and 32 indicating the different roles of the two subregions in the emotion regulations. INS1 plays a crucial role in the higher cognitive functions and participants in the processing of olfactory information.

This study, INS2 connected strongly with the ProM, Gu, PaI, STS, TP, Tha, and Amyg. The dorsal subregion Iapd, parcellated from INS2, showed higher structural connectivity with the ProM, Gu, and Tha, whereas the ventral subregion Iapv showed stronger SC with the Amyg. Previous evidence considered that the taste pathway of the macaque brain contained the nucleus of the solitary tract, thalamus, anterior insula, Gu, orbitofrontal cortex, amygdala, anterior cingulate cortex, and hypothalamus (Rolls 2019). Thus, the INS2 was mainly responsible for integrating taste information and participated in the feeding processing. A previous tracer study reported that the TP region had anatomical connectivity with agranular, parainsular, and dysgranular insula, the medial frontal, and orbitofrontal cortex, implicating the TP was associated with the auditory-related memory processing (Corcoles-Parada et al. 2019). In our study, Iapv, connected with TP and PaI, may participate in the auditory memory function.

The dorsal subregions Idd and Igd, subdivided from the INS3 and INS4 respectively, had strong connections with 6VR, verified by a tracer study (Morecraft et al. 2015), reporting the anatomical connectivity of the dysgranular insula with the ventral premotor area. In addition, Idd and Igd showed strong connections with areas 44, 45, and IPL. The previous study (Petrides and Pandya 2009) has considered that macaque areas 44 and 45 may be homologous to the human Broc’s area. The ventral premotor mainly controlled the hand (Kraskov et al. 2009) and orofacial (Ferrari et al. 2003) musculature. IPL also was demonstrated by the recent study to be homologous in the primate and was mainly involved in the tool use and language (Cheng et al. 2021). Taken together, the Idd and Igd may be a participant in the hand control and orofacial motor in the rhesus monkey, such as vocalization. In addition, the connections of Idd and Igd also showed higher connections with areas 2/1 and S2, indicating that these two subregions were mainly involved in the somatosensory functions. Concerning the ventral subregions of INS3 and INS4, Idv and Igv both had higher connections with AK and PaA, indicating that these two subregions were mainly responsible for integrating the auditory information, and together with the abovementioned subregion Iapv, thus these three ventral subregions played an vital role in the auditory function.

### 4.3 Macaque insula gradient captures structural features

After identifying the gradient of the macaque insula based on structural connectivity, we investigated the underlying architectonic mechanisms of macaque insula microstructure and morphology, discovering close correlations between insula thickness, T1w/T2w and connectivity-based gradient. The macaque insular cortex was characterized by the lamination pattern and also revealed by the architectonic gradient of cortical layers (Gallay et al. 2012a). In neuroimaging, T1w/T2w ratio, detecting the architectonic organization, is a proxy for the myelin content in the cortical regions. Moreover, the macaque insula has been considered to be an inconsistent cortical area with a differentiated layer of myeline (Mesulam and Mufson 1982; Evrard et al. 2014). Therefore, in this study, the relationship between the gradients and T1w/T2w map indicated that the spatial distribution of connectivity variation was corresponded to the myelination, suggesting that the macroscopic organization of the macaque insular cortex may capture the underlying microstructural features to some extent. In addition, we also detected the correlation between the gradient and thickness in the macaque insular cortex. This finding was consistent with the previous finding that the gradient of human thalamus correlated with its gray matter volume (Yang et al. 2020). The cortical thickness, measuring the depth of the cortical column, was the macroscopic representation of the layer physical features, which exhibited the incongruous characteristics in the macaque insula. Our finding, correlations of gradient and thickness, may indicate that the variation of structural connectivity in the macaque insular cortex was associated with its morphological structure. Overall, the relationship of gradients, T1w/T2w and thickness suggested that the macro-structural organization of the macaque insular cortex may be in accordance with its microstructure and morphology.

### 4.4 Limitations

Several limitations should be mentioned in the present study here. First, although the insula 4-subregion parcellation pattern appeared to be similar to the previous histological studies by extracting the 25% component from the global gradient, the chosen threshold was relatively subjective and required further examination. Second, due to the lack of in vivo MRI data of macaque brain collected by us, the correlation analysis on the relationship between the insula gradient and thickness was not conducted on the MRI data of the same macaque brains, which may influence our findings. Third, we did not construct the T1w/T2w map and just used the existed template to explore the relationship between the insula gradient and the T1w/T2w map, which weakened the interpretability of correlation analysis findings.

## 5. Conclusion

Using high-resolution ex vivo diffusion MRI data, this study revealed the topographic organization of macaque insular cortex globally and locally based on structural connectivity.

The gradient and parcellation of the macaque insular cortex were both similar to the histological architecture reported by the previous studies. Furthermore, permutation tests associated with connectivity fingerprint were performed to confirm the subdivisions of 8 subregions parcellated from the 4 components. Besides, the relationships between insula gradients and the T1w/T2w, thickness maps also suggested that the macroscopic connectivity variation across the cortical territory in the macaque insula may capture the underlying morphological and microstructural features. Overall, our study investigated macaque insula architecture from the perspective of structural connectivity, which may provide an insight into the following comparative research across species.

## Abbreviations

6VR: area 6 of cortex, ventral part, rostral subdivision
8AV: area 8 of cortex, anteroventral part
36R: area 36, rostral part
Acb: accumbens nucleus
AK: auditory koniocortex
Amyg: amygdaloid nucleus
Gu: gustatory cortex
INS: insular cortex
IPL: Inferior parietal lobule
MST: medial superior temporal area
PaA: paraauditory area
PaI: parainsular cortex
Pd: pallidus
PE: parietal area PE
Pir: piriform cortex
PPt: posterior parietal area
ProK: prokoniocortex
ProM: promotor
ReI: retroinsular area
ROI: region of interest
S2: secondary somatosensory cortex
Str: striatum
STS: superior temporal sulcus
Tha: thalamus
TP: temporopolar area
TPt: temporoparietal cortex

## Author contributions

Tianzi Jiang led the project. Tianzi Jiang and Lingzhong Fan were responsible for the design of the concept and study. Long Cao, Zhongchang Du and Yue Cui contributed to the writing, coding, plotting and validation of the pipeline. All the authors participated in discussions of the results and revision of the manuscript.

## Acknowledgments

This work was partially supported by Science and Technology Innovation 2030 - Brain Science and Brain-Inspired Intelligence Project of China (Grant No. 2021ZD0200200), the National Natural Science Foundation of China (Grant Nos. 31620103905, 82151307, and 82072099), the Science Frontier Program of the Chinese Academy of Sciences (Grant No. QYZDJ-SSW-SMC019), National Key R&D Program of China (Grant No. 2017YFA0105203), and the Strategic Priority Research Program of Chinese Academy of Sciences (XDB32030200).

## Conflict of interest

The authors declare that they have no conflict of interest.

## Data and code availability

The data and codes that support the findings of this study are available from the corresponding author upon reasonable request.

## Notes

### Competing Interest Statement

The authors have declared no competing interest.

