## Supplemental Figure S1, S2, Table S1 for "Structural connectivity gradient associated with a dichotomy reveals the topographic organization of the macaque insular cortex"

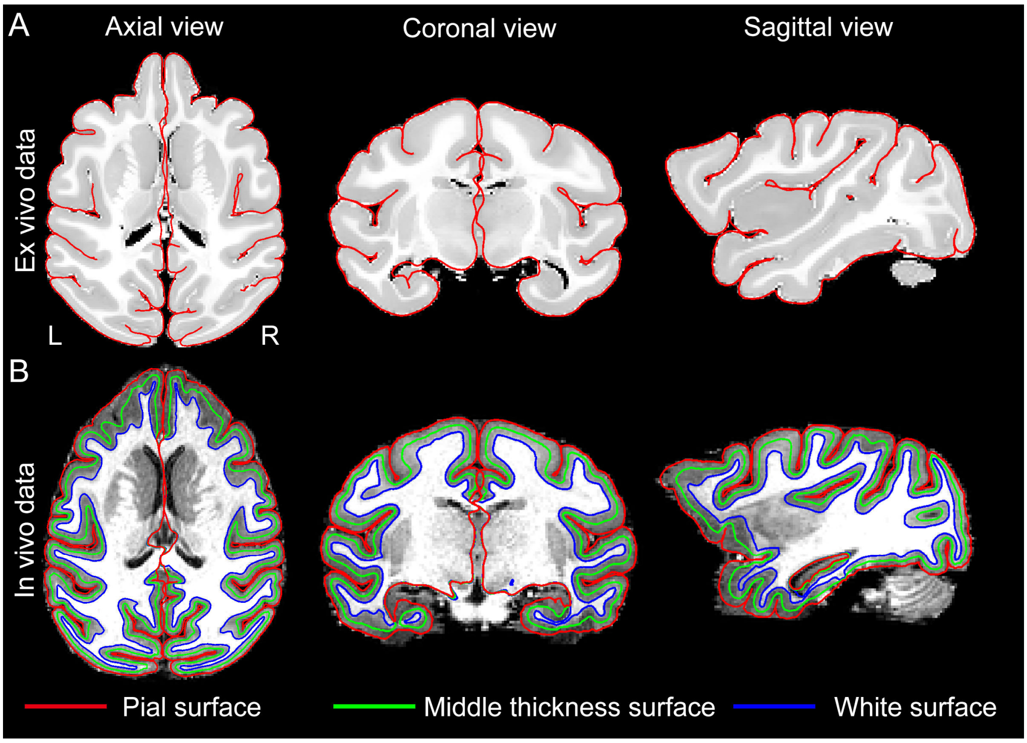


**Figure S1.** Macaque brain surface construction. (A) The pial surface of macaque ex vivo brain data. The contrast of the T2 weighted or b0 image was inverted to obtain the fake T1 image, and surface construction was mainly based on the fake T1 image. (B) The three surfaces of macaque in vivo brain data.

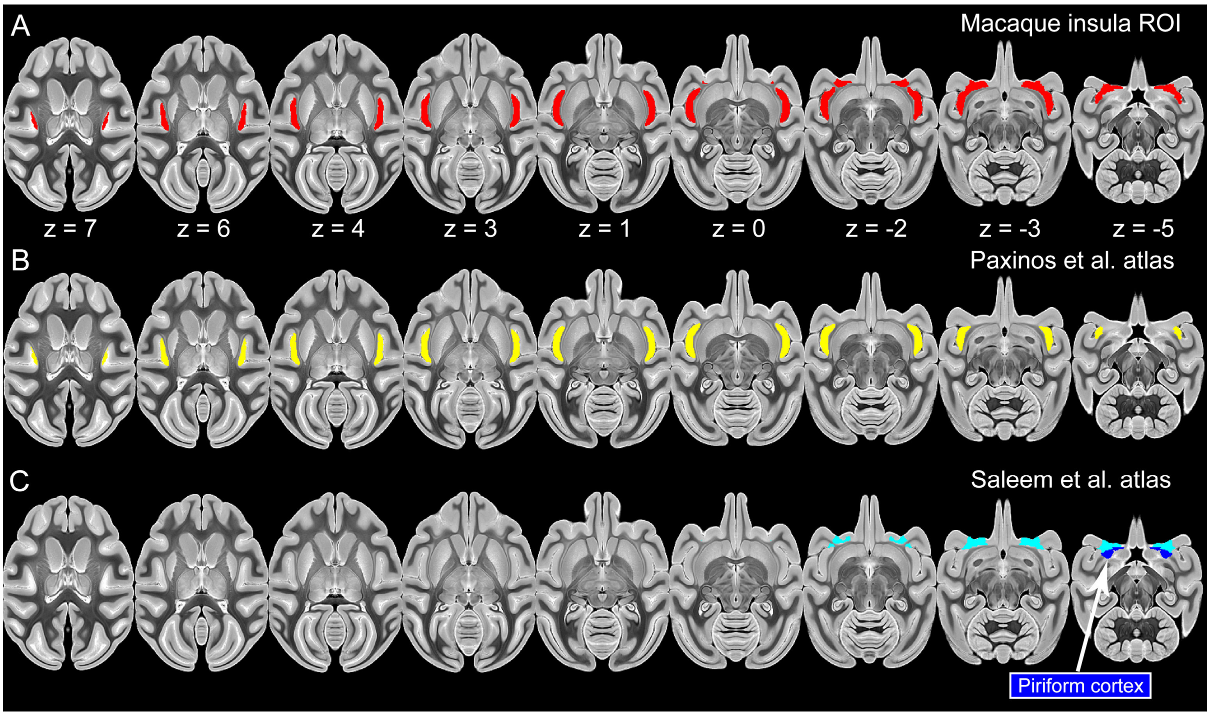


**Figure S2.** Macaque insula region of interest (ROI). (A) Macaque insula ROI was used in this study. Our insula ROI was merged from two macaque brain atlases, including Paxinos et al.'s (Paxinos et al. 2009) and Saleem et al.'s atlas (Saleem and Logothetis 2007). (B) Paxinos et al.'s macaque insula ROI. This insula ROI contained four subregions, including Ia (agranular insular cortex), Id (dysgranular insular cortex), Ig (granular insular cortex), and IPro (insular proisocortex), which were delineated by the circular sulcus and limen insula in the postmortem macaque b0 template (CIVM) (Calabrese et al. 2015b). (C) Saleem et al.'s macaque insula ROI. This insula ROI mainly occupied the posterior surface of the orbitofrontal lobe, which contained Iai, Ial, Iam, Iapl and Iapm. Given that the insula ROI was defined in the D99 space (Feng et al. 2017), nonlinear registration within Advanced Normalization Tools (ANTs) was used to warp the ROI from the D99 space to CIVM space, in which the insula ROI was adjusted by the boundary of the piriform cortex and the extended line of the inferior limiting of insula. Iai = intermediate agranular insula area; Ial = lateral agranular insula area; Iam = medial agranular insula area; Iapl = posterolateral agranular insula area; Iapm = posteromedial agranular insula area.

**Table S1**. Target regions of interest (ROIs) connected with macaque insula subregions

| ROI | Labels in the macaque atlas | ROI | Labels in the macaque atlas |
|---|---|---|---|
| area 10 | 214-217 | area 44 | 218 |
| area 45 | 219, 220 | area 46 | 221, 222 |
| area 47 | 223-225 | ProM | 226 |
| Gu | 227 | area 11 | 228-230 |
| area 13 | 231-234 | area 14 | 235, 236 |
| area 25 | 237 | 6VR | 201 |
| 8AV | 204 | area 32 | 104 |
| area 2/1 | 116 | PE | 124-126 |
| IPL | 123, 127-131 | PPt | 136 |
| S2 | 138-140 | 36R | 148 |
| AK | 149, 150 | MST | 160 |
| PaA | 162-164 | PaI | 165, 166 |
| ProK | 167-169 | ReI | 137, 170, 171 |
| STS | 172-175, 191, 192 | TPt | 193 |
| TP | 194, 195 | Tha | 6-41 |
| Pd | 44-46 | Acb | 47-49 |
| Str | 53-55 | Amyg | 63-65, 67-76, 78 |
| Pir | 77 | | |

Macaque atlas (https://www.civm.duhs.duke.edu/rhesusatlas) was delineated in the high-resolution MRI data of the postmortem rhesus macaque brain (Calabrese et al. 2015a) with Paxinos et al.'s macaque atlas (Paxinos et al. 2009). The full names of the ROIs are summarized at the beginning of the manuscript.
